# OLT1177 (Dapansutrile) inhibits Gasdermin D-dependent IL-1β Release and Pyroptotic Cell Death in Bone Marrow-derived Macrophages

**DOI:** 10.64898/2026.05.31.729061

**Authors:** Karl A. Pankratz, Masoom Raza, Jarren Ypil, Migachelle Banks, Carlo Marchetti, Tania Azam, Charles A. Dinarello, Shaikh M. Atif

**Affiliations:** University of Colorado Anschutz Medical Campus, Aurora, CO 80045

## Abstract

Gasdermins are a family of pore-forming proteins that regulate the release of pro-inflammatory cytokine, interleukin-1β (IL-1β) from infected or PAMP-stimulated cells. During infection or injury, IL-1β is released by both human and mouse macrophages. IL-1β release from mouse macrophages is associated with cell death, often termed “pyroptosis”. Mouse macrophages undergoing pyroptosis assemble an exit channel termed gasdermin D (GSDMD). Both the processing of IL-1β and the formation of the exit channel are caspase-1 dependent. Here, in bacterial endotoxin, lipopolysaccharide (LPS), treated mouse bone marrow-derived macrophages (BMDMs), we studied the pharmacologic inhibition of the intracellular nucleotide-binding domain, leucine-rich-containing family, pyrin domain– containing-3 (NLRP3) inflammasome by OLT1177. BMDMs stimulated with LPS plus the potassium efflux inducer nigericin triggered the formation of the NLRP3 inflammasome. Treatment of these BMDMs with OLT1177 suppressed cell death by 42% and ASC (apoptosis-associated speck-like protein containing a caspase recruitment domain)-speck formation by approximately 60%. In addition, OLT1177 dose-dependently inhibited IL-1β, CCL3, and myeloperoxidase (MPO) secretion and the pore-forming (GSDMD) from LPS-primed BMDMs, suggesting the existence of a vicious cycle controlled by IL-1β release. Overall, our study demonstrates that OLT1177 prevents IL-1β release from BMDMs by inhibiting caspase-1 and the conversion of (GSDMD) into its active N-terminal fragment (GSDMD-N). This study thus supports the concept that orally administered OLT1177 can be used to prevent local as well as systemic inflammation in humans.

## Introduction

Several inflammatory conditions are mediated, in part, by interleukin-1β (IL-1β); for example, gout flares ^1^, neurodegenerative diseases^2, 3, 4^ and cancer ^5^. There are two different genes termed IL-1, and both code for proteins that are expressed initially as precursors. The IL-1α precursor is primarily an intracellular cytokine, commonly found in healthy tissues such as the skin and the gastrointestinal tract, and is not inflammatory ^6^. However, when cells die, processed IL-1α is released and is highly inflammatory ^7^. In contrast, the IL-1β precursor is not normally present in the healthy state; rather, myeloid cells such as neutrophils and mononuclear phagocytes primarily produce IL-1β upon activation. Moreover, the IL-1β precursor itself is inactive and requires processing by the intracellular enzyme, caspase-1, to form an active cytokine. Once processed and released into the extracellular space, IL-1β induces local as well as systemic inflammation. For example, fever is a manifestation of IL-1β-driven systemic inflammation ^8^. There are several mechanisms by which mature IL-1β is released from living cells, including vesicles and channels ^8^. In human myeloid cells, active IL-1β release is independent of cell death, while in mouse macrophages, the release of IL-1β is associated with inflammatory cell death termed “pyroptosis”. In cells undergoing pyroptosis, several genes encoding caspases and gasdermins are increased, including S-palmitoylation of conserved cysteines (Cys191/192 resulting in cytokine release ^9^.

During the canonical NLRP3 (nucleotide-binding domain, leucine-rich-containing family, pyrin domain–containing-3) inflammasome activation, the inactive IL-1β precursor, as well as the intracellular protein gasdermin D (GSDMD), are processed by caspase-1. Processing of GSDMD results in the formation of the pore-forming N-terminal fragment (GSDMD-N) ^10^. Therefore, in addition to IL-1β activation and release, the downstream effects of the NLRP3 inflammasome play a crucial role in the formation of the GSDMD-N exit channel. In the present study, we used the small molecule sulfonyl nitrile compound OLT1177 (Dapansutrile), a known NLRP3 inhibitor, to assess pharmacologic inhibition of NLRP3 and its effects on GSDMD processing in the mouse bone marrow-derived macrophages (BMDMs). We selected OLT1177 since the compound has reduced cytotoxicity and has been administered orally in clinical studies to patients with gout ^1^ as well as heart failure ^11^, and has demonstrated benefit by attenuating both the inflammatory pain and reducing active IL-1β levels in this disease. In summary, our data showed that OLT1177 reduces cell death, cytokines, and myeloperoxidase (MPO) in LPS plus nigericin-treated BMDMs, and prevents IL-1β release and GSDMD cleavage to GSDMD-N.

## Methods

### Isolation and culture of bone marrow-derived macrophages (BMDMs)

C57BL/6 male mice (Jackson Laboratory, Bar Harbor, ME) were used in these studies. 12 to 16-week-old male and female mice were euthanized, and the femurs were removed and flushed with saline to obtain bone marrow. The bone marrow from two femurs was pooled, centrifuged at 650*g* for 10 minutes, and the cell pellet was washed twice (2X) in 0.9% NaCl before resuspending in complete media (Roswell Park Memorial Institute (RPMI)-containing 10% fetal bovine serum (FBS), referred to as RPMI-10. 1×10^5^ cells/well were seeded in a 96-well flat-bottom plate in a final volume of 200 µL, and the cells were cultured in the presence of 10% L929 (L cell, L-929, derivative of Strain L) supernatant to induce macrophage polarization ^12,13^. After 4 days at 37°C, the medium was removed, and 200 µL of fresh L929 supplemented RPMI-10 media was added for a total of 7 days.

### NLRP3 inflammasome activation and Enzyme-linked Immunosorbent assay, ELISA

On day 7, the LPS from *E. coli* OB55:B5 (Sigma-Aldrich, St. Louis, MO) was added to BMDMs in fresh RPMI-10 media at a concentration of 1 µg/mL for a total of 4 hours to prime the NLRP3 inflammasome components. After 3.5 hours, the NLRP3 inflammasome formation was induced with 10 µM nigericin (InvivoGen, San Diego, CA) for 30 minutes. To test the inhibitory potential of OLT1177, 10 minutes before adding nigericin, increasing concentrations of OLT1177 (0.1, 1, and 10 µM) were added to the cells. After 4 hours, the mouse IL-1β, CCL3 (macrophage-inflammatory protein-1 alpha, MIP-1α), myeloperoxidase (MPO), C-X-C motif chemokine ligand 1 (CXCL1), and tumor necrosis factor-alpha (TNF-α) were measured in the culture supernatants by ELISA (R&D Systems, Minneapolis, MN).

### LDH Assay

Percent cytotoxicity was examined using the LDH Cytotoxicity Assay Kit (Cayman Chemical, Ann Arbor, MI). 1×10^5^ BMDMs were cultured in a 96-well plate and treated with LPS alone or LPS/nigericin. BMDMs stimulated with LPS/nigericin were treated with various dosages (0.1-10 μM) of OLT1177. At 4h, 100 μL of the culture supernatants were transferred to the new plate, and 100 μL of the LDH reaction solution was added to each well. The plate was then incubated for 30 minutes at 37ºC. Absorbance was read at 490 nm with a plate reader (BioTek, Winooski, VT).

### Western Blotting

BMDMs were lysed in a buffer containing 50 mM Tris-HCl, 150 mM NaCl, 1.0% NP-40, 0.5% sodium deoxycholate, 1.0 mM EDTA, 0.1% SDS, and 0.01% sodium azide supplemented with a mixture of protease inhibitors (Sigma-Aldrich). The lysate was then centrifuged at 13,000*g* for 20 minutes at 4°C, and the supernatants were collected and stored at −80 °C. Protein concentration was determined in the clarified supernatant using the Bio-Rad protein assay (Bio-Rad Laboratories, Hercules, CA). Proteins were electrophoresed on Mini-PROTEAN TGX 4-20% gels (Bio-Rad Laboratories) and transferred to 0.1 μm nitrocellulose (GE Water & Process Technologies). Membranes were blocked in 5% dried non-fat milk in phosphate-buffered saline (PBS)-, 0.5% Tween for 1 hour at room temperature. Primary antibodies for GSDMD-N, IL-1β, and caspase-1 were from Cell Signaling (Danvers, MA); NLRP3 antibodies were purchased from AdipoGen, San Diego, CA. A primary antibody against β-actin (Santa Cruz Biotechnology, Santa Cruz, CA) was used to assess protein loading.

### *ASC speck* formation

ASC (apoptosis-associated speck-like protein containing a caspase recruitment domain) forms specks in BMDMs. BMDMs were treated as described above. The cells were washed with PBS, fixed with 70%-30% acetone-methanol, and incubated in a humidified slide chamber for 1 hour. 10% normal donkey serum (Jackson Immunologicals, West Grove, PA) in PBS was added to block non-specific binding sites. After removal of the donkey serum, the primary antibody to ASC (D2W8U from Cell Signaling) or isomolar, species-specific anti-IgG (R&D Systems) was added and incubated overnight at 4°C on a rotating platform. The slides were then washed three times with 1% bovine serum albumin (BSA) in PBS, and cells were incubated for 1 hour at room temperature with donkey anti–rabbit– Alexa488 (Life Technologies, Carlsbad, CA) conjugated secondary antibody. Nuclei were stained with diamidino-2-phenylindole (DAPI) (Life Technologies), and ASC specks were quantified by measuring the number of intracellular bodies with fluorescence intensity greater than 10,000 Active-Fluorescent Units (AFU) by fluorescence microscopy. Images were acquired using a Marianas Imaging Station (Intelligent Imaging Innovations, Denver, CO) using a Zeiss 639 Plan-Apochromat objective (1.4 N.A.), a Sutter Xenon light source, and a Cooke SensiCam (The Cooke Corporation, Eliot, ME).

## Results

### Reduced cell death and ASC-speck formation by OLT1177 in LPS/nigericin-stimulated BMDMs

Previously, OLT1177 showed inhibition of cell death in J774 A.1 mouse macrophage cell line and primary neutrophils^14^. Here, we have used mouse bone marrow-derived macrophages and examined the effect of OLT1177 on the cell death of BMDMs treated with LPS plus nigericin (LPS/nigericin). As shown in Figure 1A, OLT1177 significantly reduced the percent cytotoxicity in BMDMs stimulated with LPS/nigericin. A significant increase in BMDM cell death, as measured by lactate dehydrogenase (LDH) levels in the supernatants, was observed when cells were treated with LPS and nigericin. At a concentration of 0.1 µM, OLT1177 reduced the cell death by greater than 50%. This reduction in cytotoxicity is further increased to 72% with increasing concentration of OLT1177 (10 µM).

**Figure 1:**
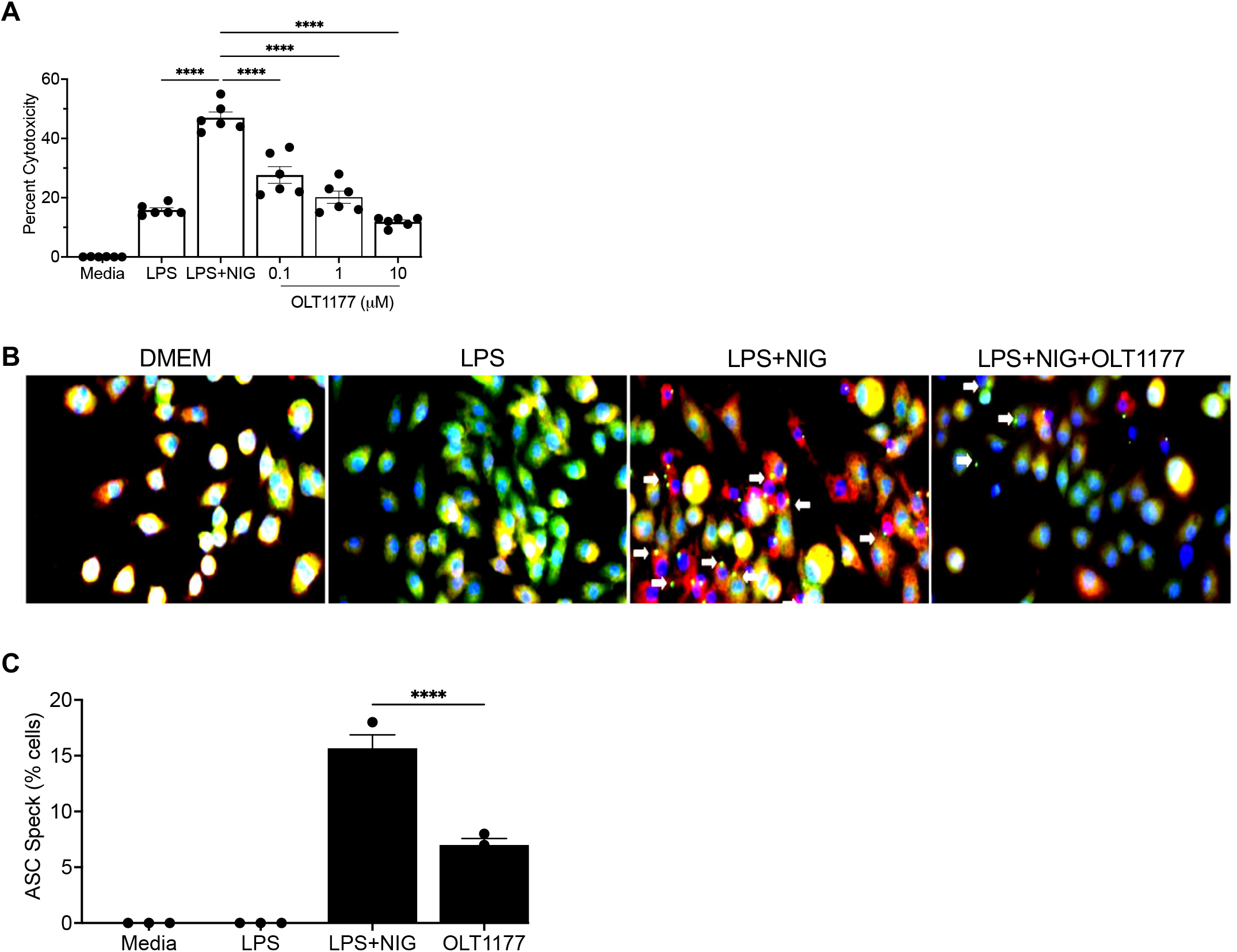
OLT1177, an NLRP3 inflammasome inhibitor, protects against LPS and Nigericin-induced cell death and ASC speck formation. **A**) BMDM from WT mice were prepared as described in Materials and Methods and treated with Media alone, lipopolysaccharide, LPS (1 μg/mL), or LPS plus nigericin (NIG, 10 μM)). Nigericin was added 30 minutes before the collection of the supernatants. The inhibitory potential of OLT1177 was tested at various dosages (0.1-10 μM) against LPS + nigericin-treated BMDMs. **B**) Immunofluorescent staining for ASC (green) and DAPI (blue) in BMDM cells stimulated with LPS and NIG in the presence of OLT1177 (10 μM). Representative ASC speck-like structures are indicated (white arrows). **C**) Mean ± SEM of ASC specks from (left-right). Data are representative of two independent biological replicates (BMDMs derived from two independent mice). ****P < 0.001.

We next examined the effect of OLT1177 on ASC speck formation in the BMDMs. ASC specks are aggregates that indicate the activation of the NLRP3 inflammasome and pyroptosis in immune cells^15^,^16^. As expected, stimulation of BMDMs with LPS did not result in the formation of ASC specks (Figure 2B). However, in the presence of LPS and nigericin, ASC specks are abundant and shown with white arrows in Figure 1B. In contrast, the number of ASC specks was significantly reduced in the presence of 10 µM OLT1177 (Figure 1B, right). Figure 1C shows the mean percent of specks in BMDM cultures treated with LPS/nigericin. As shown, the number of specks is 15% but reduced to 7% in the presence of 10 µM OLT1177 (p<0.001). Overall data suggest that OLT1177 is effective in reducing inflammation-induced cytotoxicity and ASC-speck formation.

**Figure. 2:**
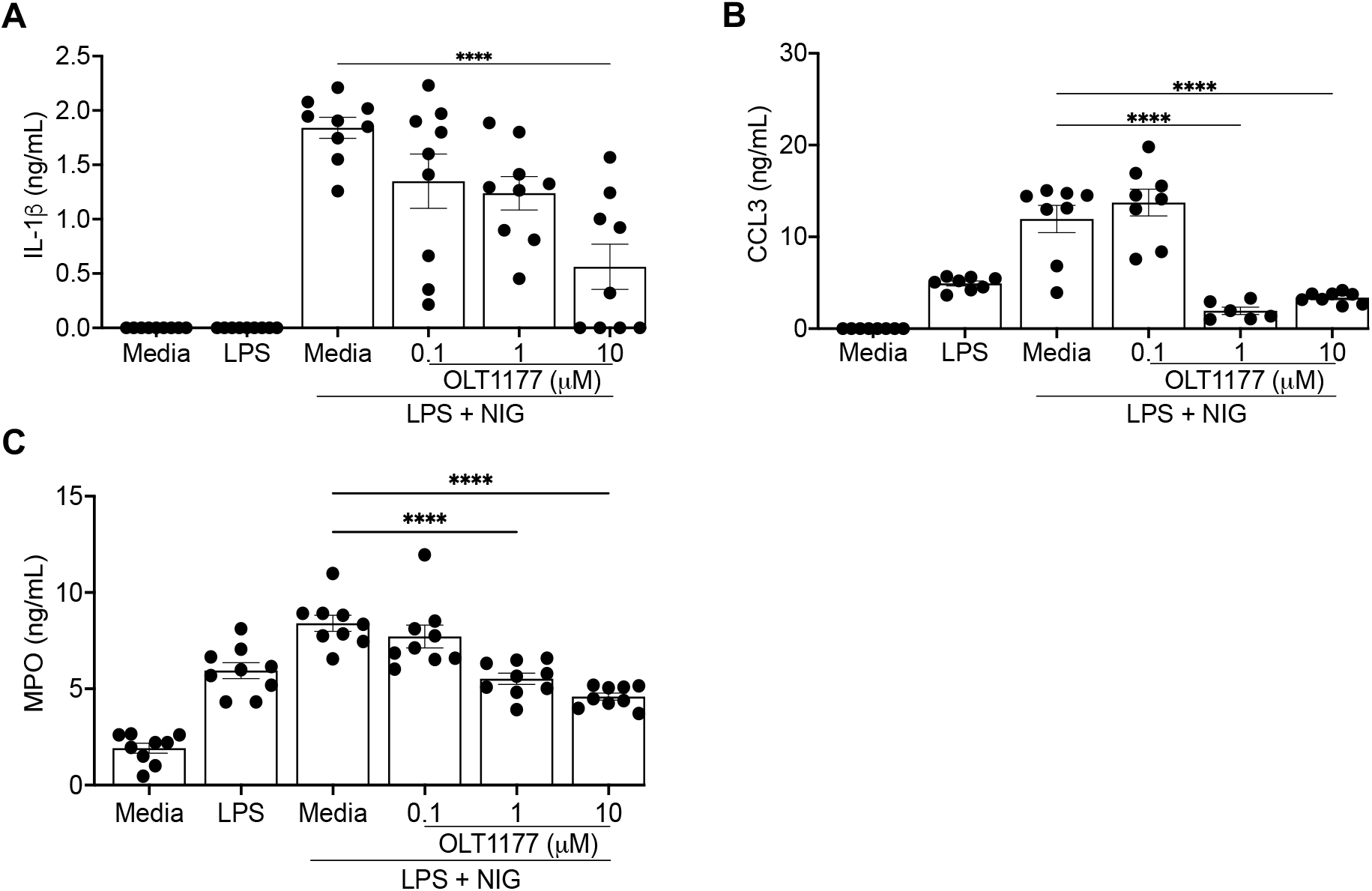
Cytokines examined in the culture supernatant of treated BMDMs. BMDMs from WT mice were treated with either media alone, lipopolysaccharide, LPS alone, LPS plus nigericin (LPS + NIG), or LPS + NIG plus OLT1177 (NLRP3 inhibitor). Nigericin (10 μM) was added to BMDMs after 3.5 hours of stimulation and cytokines were measured in the culture supernatants at 4h. IL-1β (A), CCL3 (B), and MPO (C) were measured in the culture supernatants by the R&D Duo set ELISA kit. Data are presented as mean ± SEM from three independent biological replicates (BMDMs derived from three independent mice). ****P< .0001.

### OLT1177 suppresses proinflammatory cytokines and MPO secretion from LPS/nigericin-stimulated BMDMs

Given the potential protective effect of OLT1177 against LPS/nigericin-induced cytotoxicity, we subsequently investigated whether OLT1177 influences the secretion of proinflammatory cytokines by BMDMs. In the LPS/nigericin-treated BMDMs, OLT1177 inhibited the release of IL-1β, CCL3, and MPO in a dose-dependent manner (Figure 2A-C). OLT1177 has little or no effect on TNF-α secretion from LPS plus nigericin (LPS+NIG)-stimulated macrophages. At the highest concentration, 10 µM, OLT1177 caused a significant inhibition in IL-1β release (~75%) in BMDM, whereas a 100-fold lower concentration (0.1µM) inhibited 35% (Figure 2A). Besides IL-1β, we observed a similar effect of OLT1177 on CCL3 and MPO secretion. However, no significant differences were observed in CXCL1 and TNF-α expression in the LPS+NIG treated groups in the presence or absence of OLT1177 (Supplementary Figure 1). These data suggest that OLT1177 effectively blocks the secretion of IL-1β and CCL3 but not all the cytokines from BMDM.

### Activation of caspase-1

Processing of the inactive IL-1β precursor to active IL-1β requires active caspase-1^17^. Therefore, we examined the effect of OLT1177 on the processing of pro-caspase-1 to active caspase-1. The inactive caspase-1 precursor is processed into a 22 kDa active molecule. Caspase-1 is autocatalytic and is increased in BMDMs treated with LPS+NIG ^18^. In Figure 3A, the western blot shows a progressive reduction of the 22 kDa caspase-1 protein expression with increasing concentrations of OLT1177. Figure 3B represents the dose-dependent effect of OLT1177 on the relative density of caspase-1 relative to β-actin levels in the same cells (Figure 3A). As shown, OLT1177 at 10 µM, 1 µM, and 0.1 µM reduced the 22 kDa caspase-1 by 61%, 34%, and 28%, respectively.

**Figure 3.**
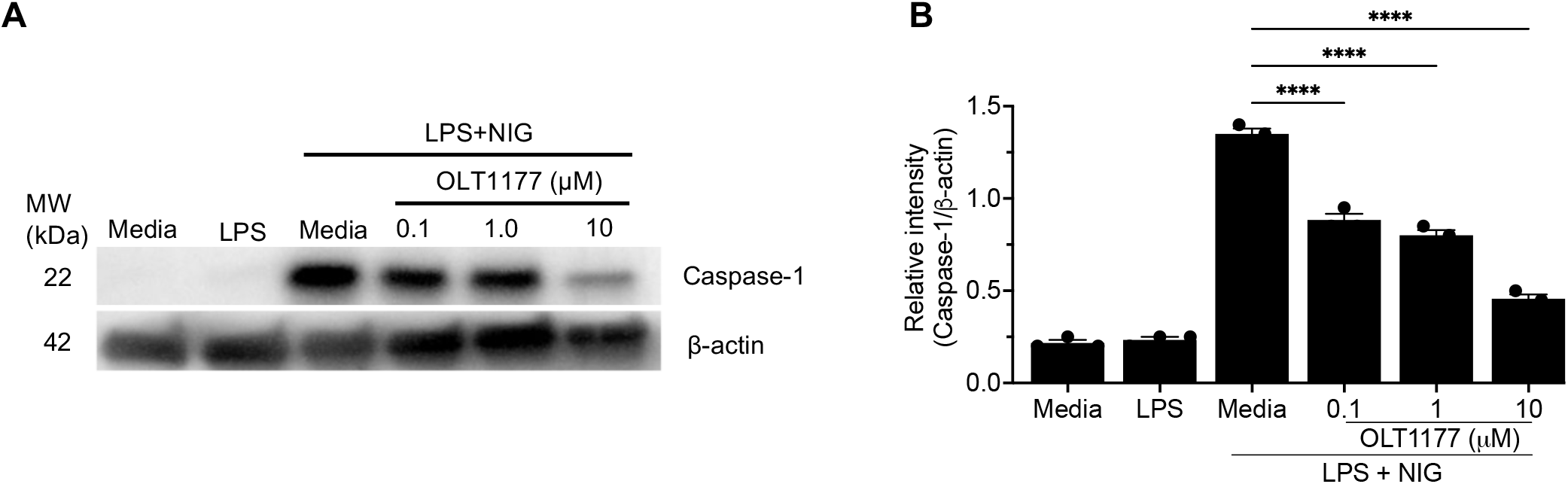
OLT1177 inhibits caspase-1 expression in LPS/nigericin-treated BMDMs. A) BMDMs were stimulated with lipopolysaccharide, LPS (1μg/mL) alone or LPS plus nigercin (NIG,10 μM). Nigericin was added to the wells after 3.5h of stimulation. OLT1177 was added to the wells, 10 minutes before the addition of nigericin and caspase-1 protein expression was determined by western blotting. All concentrations of OLT1177 significantly reduce the amount of caspase-1 in a dose-dependent manner in LPS/nigericin-treated BMDMs. B) A representative densitometric analysis of the western blot bands of caspase-1 from A. Data are representative of three independent biological replicates (BMDMs derived from three independent mice). ****P< 0.0001.

### OLT1177 treatment inhibits GSDMD processing

We next examined the effect of increasing concentrations of OLT1177 on the cleavage of pore-forming protein GSDMD. GSDMD pores facilitate the release of mature IL-1β in pyroptosis and are formed by a caspase-1-dependent cleavage of GSDMD to GSDMD-N ^10^. As shown in Figure 4A, western blotting of BMDM lysates depicts the 58 kDa full-length GSDMD (top row) in BMDMs primed with LPS. This priming also resulted in the expression of 37 kDa IL-1β precursor. However, it is only with the addition of nigericin that the cells produce the pore-forming 34 kDa GSDMD-N because the addition of nigericin induces caspase-1 expression (Figure 3).

**Figure 4.**
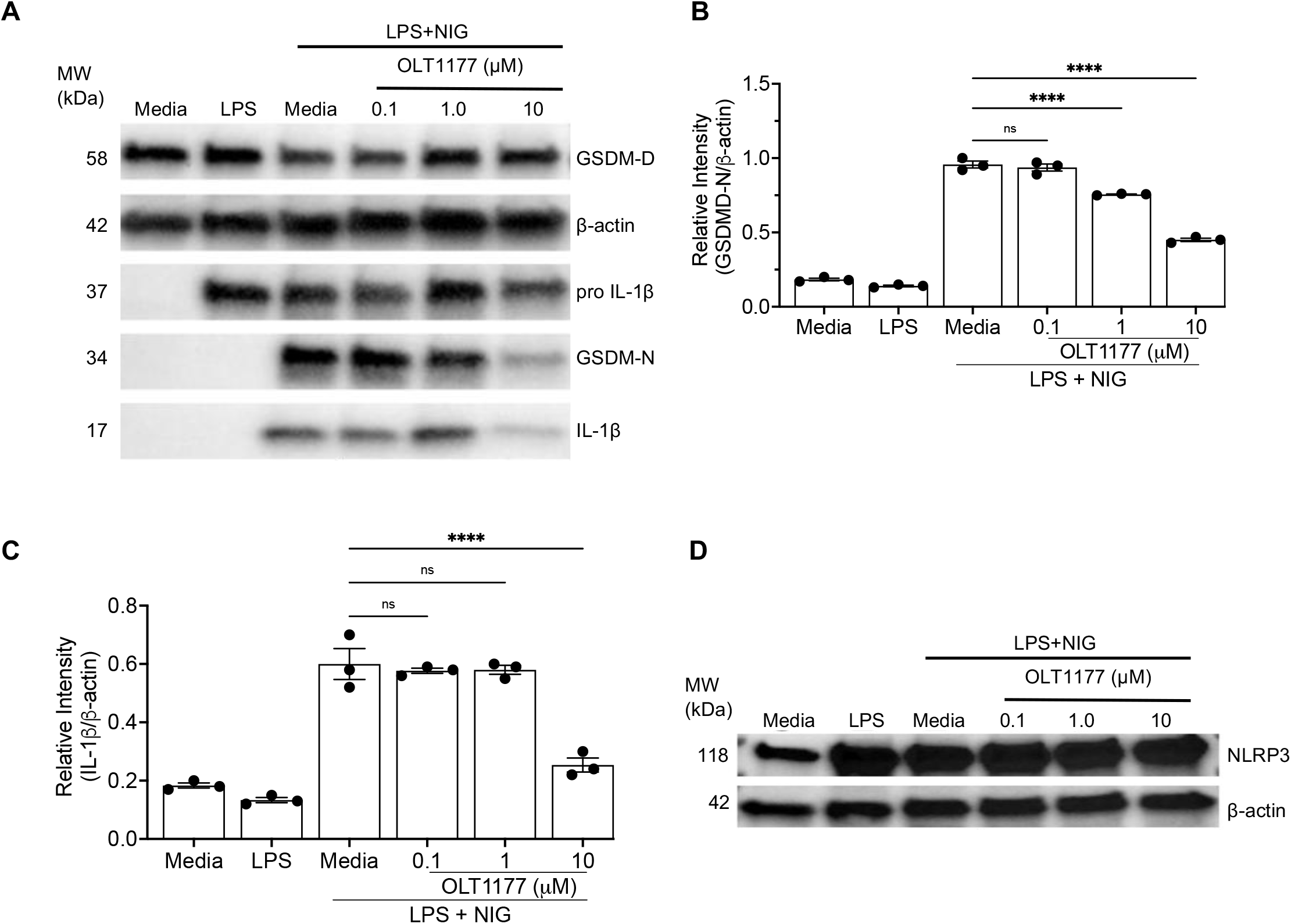
OLT1177 inhibits gasdermin, GSDMD cleavage to GSDMD-N in LPS/nigericin-treated BMDMs. A) BMDMs were treated with lipopolysaccharide, LPS (1μg/mL) and LPS plus nigercin (NIG). Representative western blots show OLT1177 inhibits GSDMD cleavage to GSDMD-N in a dose-dependent manner. (B, C) Densitometric analysis of the GSDMD-N and IL-1β bands from Figure 1A. All concentrations of OLT1177 significantly reduce both GSDMD-N and IL-1β in a dose-dependent manner. D) Western blot shows NLRP3 and β-actin expression in BMDMs. Data are representative of three independent biological replicates (BMDMs derived from three independent mice). ****P< 0.0001.

When BMDMs were treated with increasing concentrations of OLT1177, there was less GSDMD-N, which was quantified in Figure 4B. The western blot and quantification (Fig. 4A and Fig. 4C) reveal that at 10 µM, OLT1177 reduces IL-1β by 58%. The decrease in GSDMD-N at 10 µM OLT1177 was consistent with the levels of IL-1β in the supernatants shown in Figure 2A. Also shown in Figure 4A is the reduction in the active 17 kDa IL-1β, which is quantified in Figure 4C. The reduction in IL-1β in Figure 4, examined by western blotting, is comparable to the quantification by the ELISA method (Figure 2). There was no reduction in NLRP3 protein expression in the presence of OLT1177 (Figure 4D).

## Discussion

OLT1177 is a selective inhibitor of the NLRP3 inflammasome that has been administered as an oral therapeutic to subjects with acute gout flares, a prototypical disease of NLRP3 activation ^19^. OLT1177 attenuated inflammatory joint pain mediated by IL-1β activation and release, known to be triggered by monosodium uric acid (MSU) crystal deposits in the synovial space. In this study, we elucidated the downstream effects of NLRP3 inhibition by OLT1177. Firstly, OLT1177 reduced the *in vitro* release of processed IL-1β in mouse BMDMs by preventing the conversion of GSDMD to GSDMD-N. Although the release of IL-1β from human monocytes has been documented ^8, 10^, most studies that focus on the mechanisms of IL-1β release have been conducted in mouse cells, particularly in mouse BMDMs. Here we focused on two areas: blocking the maturation of IL-1β and inhibition of cytokines for reducing inflammation by OLT1177 and rescuing from pyroptotic cell death.

We and others previously have reported the effect of OLT1177 on the mechanism of potassium efflux upon activation of the pentraxin receptor (P2X7) by ATP ^14^. In that study, ion currents were measured in whole-cell patch-clamp methods in human LPS-primed monocytic U937 cells. An increase in ion currents was observed following activation of the P2X7 receptor; however, OLT1177 did not alter the currents. We conclude that OLT1177 suppression of IL-1β release was independent of ion-mediated current changes. As expected, OLT1177 inhibited the formation of the NLRP3 inflammasome in cultured mouse macrophages with the reduction in ASC speck formation (Figure 1). ASCs are critical for the NLRP3 inflammasome assembly and activation, and inflammasome-dependent speck formation (Figure 1B). As shown in Figure 1C, the number of specks was significantly decreased with 10 µM OLT1177. Relevant to this discussion, in the resting state, OLT1177 binds to recombinant NLRP3 with a K_d_ of 1.2µM ^20^. However, *in vitro* as well as *in vivo*, OLT1177 binds to natural NLRP3, which accounts for the reduction in molecules, including CCL3, MPO, involved in inflammation. CXCL1 and TNF-α were not affected by OLT1177 treatment, suggesting that these molecules are regulated independently of NLRP3 inflammasomes. Both CXCL1 and TNF-α were shown to play an important role in inflammation by causing the recruitment and activation of immune cells^21–23^.

Active caspase-1 has two major roles for IL-1β-mediated inflammation; the primary function is the processing of the inactive IL-1β precursor into active IL-1β, and the second function is the processing of GSDMD to form GSDMD-N exit channels. This dual function of caspase-1 is an example of the economy of NLRP3 activation. As revealed in the western blot (Figure 3A and B), the activation by LPS and nigericin increases caspase-1, which is dose-dependently inhibited by OLT1177. We also observed that unprocessed GSDMD is expressed in BMDMs from healthy mice (Figure 4A, top row).

Nevertheless, as shown in Figure 4D, NLRP3 remains unchanged, even when BMDMs are stimulated with LPS plus nigericin. With the cleavage of procaspase-1, active caspase-1 functions as a mature 22 kDa molecule (Figure 3), resulting in the GSDM-N pores^17, 24, 25^. In mouse BMDMs, the exit of IL-1β is linked to cell death (Figure 1) and hence the use of the term pyroptosis^26, 27^. In peripheral blood mononuclear cells from humans with acute gout flares, one can observe the intracellular processing of the IL-1β precursor as well as the reduction of the processing by OLT1177^19^. In mouse BMDMs, LPS plus nigericin results in the accumulation of GSDMD-N and 17 kDa IL-1β, whereas LPS alone only induces the IL-1β precursor. The comparison between LPS only and LPS plus nigericin is in the formation of the GSDMD-N pores, which allows IL-1β to exit the cells through the pore inserted into the phospholipid bilayer of the cell.

In Figure 5, we summarized our findings and highlighted the role of caspase-1 as the centerpiece of these studies. As shown, the processing of the IL-1β precursor and the formation of the GSDM-N exit pore occur as caspase-1 becomes active. Also shown in Figure 5, there is a direct correlation between activation of NLRP3 and caspase-1 activity. Although we illustrate LPS as the activator of the IL-1β precursor, several stimulators result in a cascade that leads to the synthesis of the IL-1β precursor. Overall, this study highlights the important role of OLT1177 in blocking the IL-1β release and GSDM-D processing to GSDM-N. Additionally, in BMDMS, OLT1177 dose-dependently reduced cellular toxicity by manyfold.

**Figure 5.**
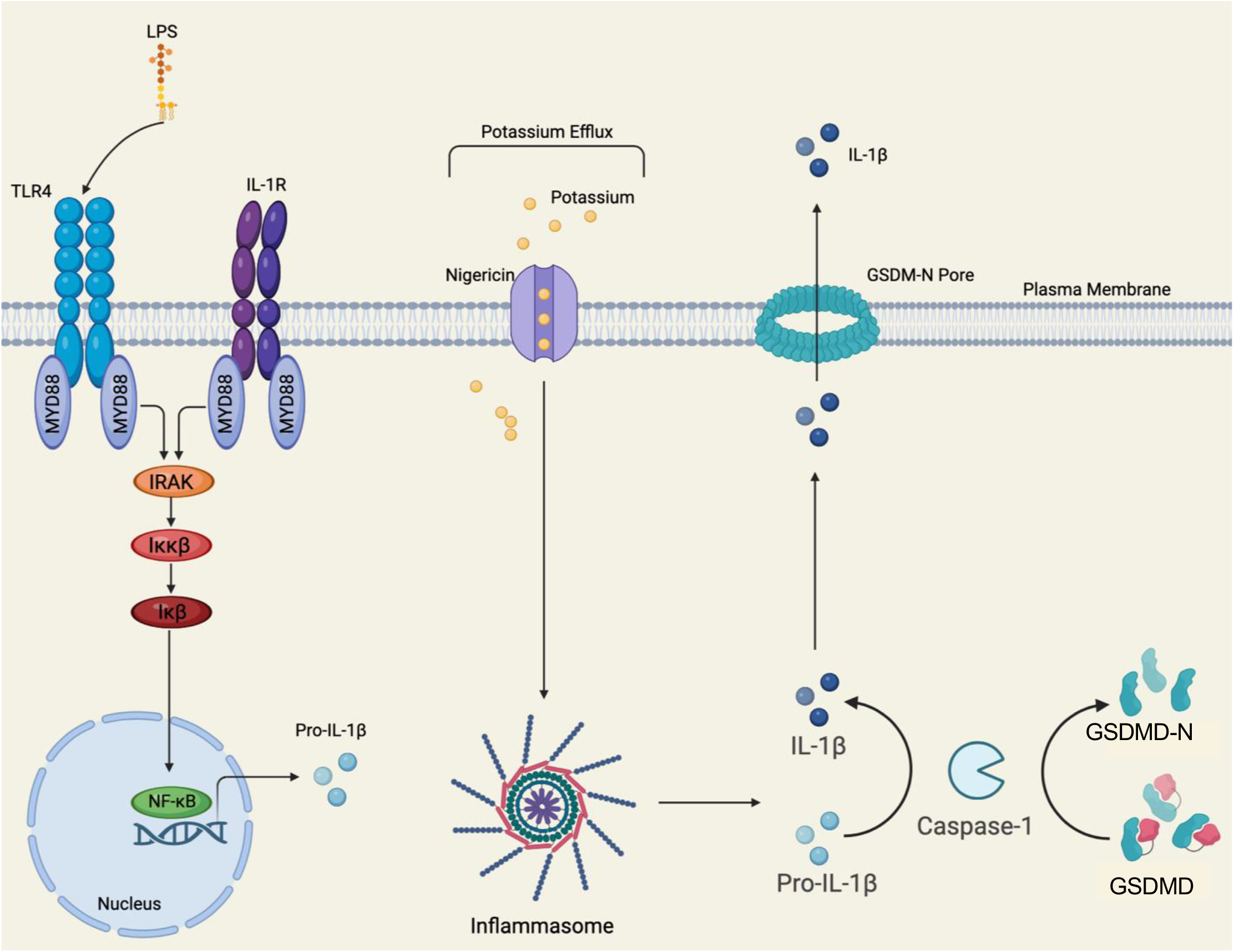
Schematics of IL-1β release via GSDM-D/GSDM-N pores in LPS/nigericin-stimulated macrophages. LPS induces cell activation and expression of the NLRP3 gene. Nigericin is a polyether antibiotic and acts as a potassium ionophore, causing NLRP3 inflammasome activation. Caspase-1 cleaves pro-IL-1β and GSDMD to IL-1β and GSDM-N, respectively. Mature IL-1β then exits the cells through the GSDMD-N pores.

## Acknowledgements

These studies are supported by NIH Grant AI 15614 (to CAD) and the Interleukin Foundation (to KAP), NIH Grant HL152576, and TRSP award (to SMA). Olatec, LLC provided OLT1177 for these experiments.

## Notes

Conflicts of Interest: K.A.P., no conflicts; M.R., no conflicts; J.Y., no conflicts, M.B., no conflicts; T. A., no conflicts; C.A.D. serves as Chairman of Olatec’s Scientific Advisory Board, is co-Chief Scientific Officer, receives compensation, and has equity in Olatec; C.M. serves as Director for Olatec’s Innovative Science Program and has equity in Olatec. S.M.A., no conflicts.

### Competing Interest Statement

A conflict of interest statement is mentioned in the manuscript.

