## Supplementary Figure 1 for "OLT1177 (Dapansutrile) inhibits Gasdermin D-dependent IL-1β Release and Pyroptotic Cell Death in Bone Marrow-derived Macrophages"

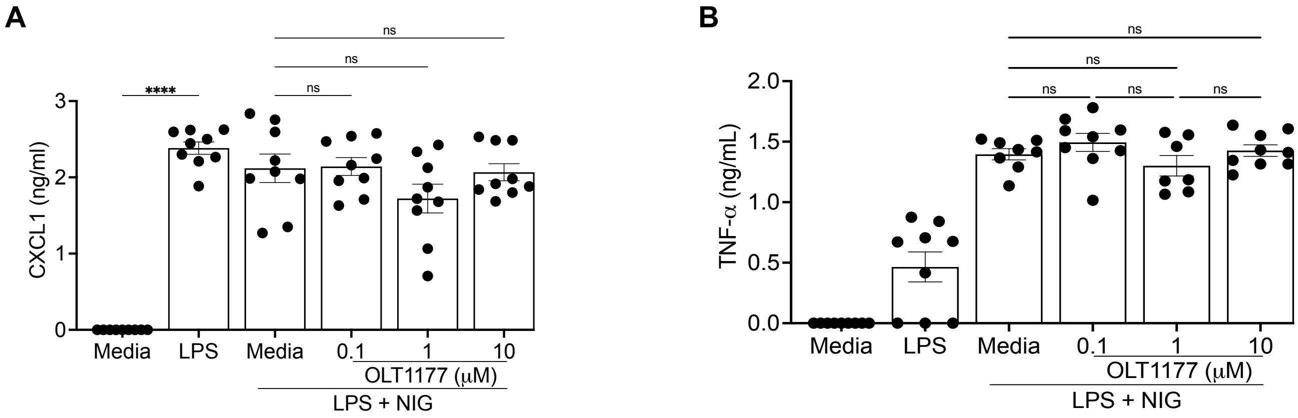


**Supplementary Figure. 1: CXCL1 and TNF-α are not inhibited by OLT1177 treatment in LPS plus nigericin-treated BMDMs**. BMDMs from WT mice were treated with either media alone, LPS, LPS plus nigericin (LPS + NIG), or LPS + NIG plus OLT1177 (NLRP3 inhibitor). Nigericin (10 μM) was added to BMDMs after 3.5 hours of stimulation and cytokines were measured in the culture supernatants at 4h. CXCL1 and TNF-α were measured in the culture supernatants by the R&D Duo set ELISA kit. Data are presented as mean ± SEM from three independent biological replicates (BMDMs derived from three independent mice).****P<0.0001.
